# Synchrony within Aβ mechanoreceptor subtypes governs signal propagation to primary somatosensory cortex

**DOI:** 10.64898/2026.01.16.700009

**Authors:** Wanyi Liu, Andrew E. Worthy, Alan J. Emanuel

## Abstract

Fast-conducting Aβ slowly adapting type I low threshold mechanoreceptors (SAI-LTMRs) and Aβ rapidly adapting type I (RAI-) LTMRs are critical for discriminative touch from glabrous skin. Recent studies used mouse genetic loss-of-function and optogenetic gain-of-function manipulations to determine that signals from these Aβ LTMRs are integrated subcortically to build the cortical representation of touch. However, the precise influence of each subtype on downstream responses is unclear, in part because previous manipulations lacked the capacity to reproduce physiological firing patterns in individual Aβ LTMR subtypes. Here, we took advantage of a fast variant of channelrhodopsin, CatCh, which enabled us to generate physiological spiking rates and patterns in either subtype while monitoring responses in mouse primary somatosensory cortex (S1). Doing so revealed that propagation of steady-state signals from sustained responses of Aβ SAI-LTMRs to neural activity (spiking and local field potential) in S1 is dependent on synchronous activation of multiple Aβ SAI-LTMRs. Asynchronous activation of the same Aβ SAI-LTMRs rarely produced sustained responses in S1 measured at the level of single unit spiking as well as local field potentials. This suggests that the irregular firing patterns of Aβ SAI-LTMRs during static indentations contribute to the preponderance of transient cortical responses. By contrast, both synchronous and asynchronous activation of multiple Aβ RAI-LTMRs resulted in robust S1 responses. Overall, the temporal patterning and synchrony of activity within Aβ LTMR subtypes govern how well their signals propagate through the ascending somatosensory system.

**Key Points:**

- CatCh, a sensitive and fast channelrhodopsin variant, enables pulsed-light generation of physiological spiking patterns in Aβ low threshold mechanoreceptor (LTMR) subtypes.
- The synchrony of multiple mechanoreceptors within a subtype controls the extent to which their signals influence activity in primary somatosensory cortex.
- Aβ rapidly adapting type I (RAI-) LTMRs drive stronger cortical responses than Aβ slowly adapting type I (SAI-) LTMRs, especially when considering asynchronous activation of Aβ LTMRs within each subtype.

## Introduction

Our sense of touch is one of the primary ways we interact with the external world (Reed *et al*., 1996; de Haan & Dijkerman, 2020). Physical stimuli are transduced into electrical impulses by a diverse array of mechanically sensitive peripheral nerve endings (Handler & Ginty, 2021). In mammalian glabrous (non-hairy) skin, Aβ low-threshold mechanoreceptors (LTMRs) are the main transducers of gentle and discriminative touch (Johnson, 2001; Walcher *et al*., 2018). Two major subtypes of Aβ LTMRs are the Aβ rapidly adapting type I (RAI)-LTMRs that innervate Meissner corpuscles and the Aβ slowly adapting type I (SAI)-LTMRs that are associated with Merkel cells (Johnson, 2001; Handler & Ginty, 2021). Aβ RAI-LTMRs produce action potentials at the onset and offset of ramp-and-hold mechanical stimuli and are especially sensitive to moving stimuli and vibrations at frequencies between 30 and 100 Hz (Johansson & Vallbo, 1979; LaMotte & Whitehouse, 1986; Vega-Bermudez & Johnson, 1999; Neubarth *et al*., 2020). Aβ SAI-LTMRs, by contrast, produce a burst of action potentials at touch onset and maintain spiking with irregular interspike intervals during the static phase of sustained touch (Iggo & Muir, 1969; Knibestol & Vallbo, 1980; Woodbury & Koerber, 2007; Wellnitz *et al*., 2010; Maksimovic *et al*., 2014; Al-Basha & Prescott, 2019).

Sensory inputs from these Aβ LTMRs and other mechanoreceptors are relayed to the primary somatosensory cortex (S1) through the spinal cord, dorsal column nuclei, and the thalamus, where integration and transformation of the signals can occur at each stage (Abraira & Ginty, 2013; Emanuel *et al*., 2021; Suresh *et al*., 2021; Chirila *et al*., 2022). The majority of touch-responsive S1 neurons generate a transient response that resembles the ON/OFF pattern of Aβ RAI-LTMRs in response to sustained touch (Sur *et al*., 1984; Pei *et al*., 2009; Callier *et al*., 2019; Emanuel *et al*., 2021). In rodents, selective, developmental ablation of glabrous-skin Aβ RAI-LTMRs reduces the amplitude and sensitivity of the cortical response to touch but does not change its transient ON/OFF firing pattern (Emanuel *et al*., 2021). It remains unclear how inputs from only Aβ SA-LTMRs are transformed from a sustained firing pattern into this transient ON/OFF response pattern. The slow recovery time of the ReaChR opsin we previously used to activate Aβ LTMR subtypes only allowed for the generation of action potentials at low rates (< 5 Hz) and limited our ability to infer the impact of Aβ LTMR firing at physiological rates and temporal patterns (Emanuel *et al*., 2021). Therefore, we expressed an opsin with fast kinetics, CatCh (Kleinlogel *et al*., 2011), in Aβ RAI-LTMRs or Aβ SAI-LTMRs to determine how trains of action potentials evoked at physiological rates and in physiological patterns in sensory neuron subtypes influence S1 activity.

## Methods

### Mice

All experimental procedures were approved by Emory University Institutional Animal Care and Use Committee (Protocol 202200102) and were performed in compliance with the Guide for Animal Care and Use of Laboratory Animals. Mice were housed in temperature- and humidity-controlled facilities in a 12h:12h light:dark cycle. To selectively express the CatCh opsin (fused to EYFP) in Aβ SAI-LTMRs and Aβ RAI-LTMRs we generated mice with the genotypes *TrkC^CreER^;Avil^FlpO^;R26^LSL-FSF-Catch^* and *TrkB^CreER^;Avil^FlpO^;R26^LSL-FSF-Catch^*, respectively. All alleles were previously described: *TrkC^CreER^*: (Bai *et al*., 2015), JAX 030291; *TrkB^CreER^*: (Rutlin *et al*., 2014), JAX 027214; *Avil^FlpO^*: (Choi *et al*., 2020); *R26^FSF-LSL-CatCh^* [Ai80]: (Daigle *et al*., 2018); JAX 025109. To selectively induce Cre recombination in Aβ SAI-LTMRs, we administered 2-3 mg of tamoxifen (dissolved in sunflower seed oil) to the pregnant dam at embryonic day 12.5 (E12.5) via oral gavage. To selectively induce Cre recombination in Aβ RAI-LTMRs, we administered 0.5 mg tamoxifen (i.p.; dissolved in sunflower seed oil) at postnatal day 3 (P3). All mice were on mixed C57Bl/6J and CD1 backgrounds. A total of eight *TrkC^CreER^;Avil^FlpO^;R26^LSL-FSF-Catch^* mice (3 males and 5 females) and seven *TrkB^CreER^;Avil^FlpO^;R26^LSL-FSF-Catch^* mice (3 males and 4 females) were included.

### In vivo dorsal root ganglion electrophysiology

Anaesthesia was introduced with urethane (1.5 g/kg body weight) and maintained using isoflurane (1-2%). After the induction of anaesthesia, we shaved the back of the mouse and made a midline incision over the lumbar vertebrae. We applied a spinal clamp (Kopf) to the L3 vertebra to secure the spinal column. We dissected the paravertebral muscles over the L4 vertebra and removed the bone covering the L4 dorsal root ganglion (DRG) with a fine-tipped rongeur. The surface of the L4 DRG was cleaned and then submerged in HEPES-buffered saline.

A 32-channel silicon probe (ASSY-37 H7b, Cambridge NeuroTech) was inserted into the L4 DRG. We gently brushed the mouse’s hindpaw with a paintbrush or a cotton-tipped applicator to identify neurons that respond to mechanical stimulation of the plantar surface. Raw signals were digitised and amplified through an Intan RHD 32 channel headstage (Intan Technologies) and an Open Ephys acquisition board (Siegle *et al*., 2017). Data acquisition was controlled with Open Ephys Software (version 0.6.7).

For optical stimulation of Aβ LTMRs expressing CatCh, pulses of light were generated by a 300 mW, 445-nm laser (CST-H-445-300, Ultralasers). Light was directed to different locations on the paw through two scanning galvanometer mirrors (GVS002, Thorlabs) and an Fθ lens (FTH160-1064-M39, Thorlabs). The intensity of the laser measured at the skin was 20-22 mW and the beam was focused to a spot size of 5,105 μm^2^. To measure the spot size, we attenuated the laser with a 4.0 neutral density filter and illuminated a lens-less CMOS sensor with an exposure time of 5 μs, which prevented saturation. Area was calculated from the resulting image binarised at 1/e^2^ maximum intensity.

Laser pulses (0.3 ms in duration; 6.0-6.6 μJ) were delivered to the glabrous skin region of the hindpaw at each point within an 8 x 6 mm area (0.2 mm spacing) to map optical receptive fields of CatCh-expressing neurons. To assess how reliably action potentials follow the optical stimulation, we delivered two laser stimulus sets to the receptive field of that neuron: (1) a series of increasing frequencies ranging from 2 to 40 Hz, and (2) the irregular temporal pattern used for the synchronous stimulation (10 pulses in 0.5 s; see below).

After the DRG recording, mice were euthanised with cervical dislocation. The brain and paws were extracted and preserved with post-fixation overnight (4% paraformaldehyde for brain and Zamboni fixative for paws). The tissue samples were then washed with PBS, cryoprotected with 30% sucrose, and frozen embedded in optimal cutting temperature (OCT) compound.

### Immunohistochemistry

We cryosectioned glabrous skin pedal pads into 25 μm sections using a cryostat (Leica) and collected sections on glass slides (Fisher 1255015). We stained sections by washing twice for 5 min each with 1xPBS containing 0.1% Triton X-100 (0.1% PBST), then incubating with blocking solution (5% normal donkey serum [Jackson ImmunoResearch Labs 017-000-121] in 0.3% PBST) for 1h at room temperature, and then incubating with primary antibodies in blocking solutions at 4 °C overnight. Primary antibodies included chicken polyclonal anti-green fluorescent protein (Aves Labs GFP-1020; 1:500), rabbit polyclonal anti-S100 β (Thermo Fisher 15146-1-AP; 1:500), and rat monoclonal anti-Troma1 (DSHB AB_531826; 1:200). We then then washed the tissue four times (15 min) with 0.1% PBST and subsequently incubated with secondary antibodies (donkey anti-chicken conjugated to Alexa 488, Invitrogen A78948; donkey anti-rabbit conjugated to Alexa 546, Invitrogen A10040; donkey anti-rat conjugated to Alexa 647, Invitrogen A48272; all 1:500) in blocking solution for 2 h at room temperature. We washed three times (15 min) with 0.1% PBST and included DAPI solution (5 μg/mL; AAT Bioquest) in the second wash. Finally, we washed with 1x PBS for 15 min and mounted with Fluoromount-G (Southern Biotech). We imaged tissue with a Keyence BZX-7000 automated fluorescence microscope.

### Surgery and craniotomy

We administered dexamethasone (2 mg/kg) to adult (>P56) mice 4-12 hours prior to surgery to minimise swelling. Immediately before surgery, we administered sustained-release meloxicam (2-4 mg/kg; Wedgewood Pharmacy). During surgery, mice were anesthetised with 1.5-2% isoflurane and locally at the incision with Lidocaine (1%, 0.03 mL). We removed the scalp and periosteum, dried the skull, and mounted a titanium headplate to the skull with dental acrylic (Metabond). We made a circular craniotomy (approximately 1 mm in diameter) above forepaw S1 (right hemisphere: 2.10 mm lateral, 0.00 mm posterior to bregma). We sealed the craniotomy with Kwik-Sil or Kwik-Cast (WPI) and cemented an aluminium ring onto the headplate to provide a well for a recording bath solution. Extended-release buprenorphine (0.5-1 mg/kg; Wedgewood Pharmacy) was administered post-operatively for analgesia. Mice recovered overnight before the first recording session.

### In vivo cortex electrophysiology

All S1 recordings were performed in awake, head-fixed mice. Prior to each recording, the mouse was habituated to head fixation for 10 minutes. We removed the craniotomy cap and submerged the craniotomy in a HEPES-buffered saline solution (150 mM NaCl, 2.5 mM KCl, and 10 mM HEPES; pH = 7.4). We inserted a Neuropixels 1.0 probe coated with DiI (D3911, Thermo Fisher) into forepaw S1 of the right hemisphere and advanced the tip of the probe to approximately 1,600 μm below the dura. To verify S1 placement, we gently brushed the mouse’s skin on the contralateral side (left) with a fine paintbrush. We monitored brush-evoked spikes from multiple recording channels to verify the approximate location of the receptive field. If the receptive field was not on the forepaw pedal pads, we removed the probe from the brain and reinserted it into a new location. We restrained the left forepaw with electrical tape and placed it over a rectangular aperture cut into an acrylic platform supporting the mouse.

We acquired data using the Neuropixels PXIe control system (IMEC) and the Open Ephys GUI software (Siegle *et al*., 2017). Raw signals were digitised, amplified, filtered, and saved into two bands: AP band: digitised at 30 kHz, amplified 500x, and bandpass filtered at 300-10,000 Hz, and LFP band: digitised at 2.5 kHz, amplified 250x, bandpass filtered at 0.5-500 Hz.

### Optical skin stimulation during S1 recordings

In the first experiment, we delivered all light stimuli (generated with the same stimulator described for DRG recordings) to the centre of the forepaw contralateral to the recording site. Stimulation patterns included regularly spaced trains of laser pulses at 5, 20, and 40 Hz over 0.5 s as well as a train of laser pulses that emulated the physiological response of a single Aβ SAI-LTMR over 0.5 s (“physiological” laser stimulation). Each laser pattern was repeated 50 times with 5-s inter-trial intervals. For this experiment, each stimulus was presented in a separate block of trials.

In a separate group of mice, we delivered the laser pulses across the 5 pads on the mouse’s paw. The x and y coordinates of the centre of the 5 pedal pads on the left forepaw were calibrated. In each trial, trains of laser pulses were delivered either synchronously or asynchronously to each of 5 locations equally spaced on a circle (0.6-mm diameter) centred on one pad. Each train included 10 laser pulses within 0.5 s. For synchronous stimulation, pulses were delivered to each of the five locations in rapid succession, with a 0.4-ms delay between adjacent pulses (the minimum interval possible given the response time of the galvanometer mirrors). For asynchronous stimulation, we delivered pulses to the five locations with inter-pulse intervals of at least 3 ms, preventing synchronous activation. For each of the 5 pads, both the synchronous stimulation and asynchronous stimulation trials were repeated for 30 trials, resulting in a total of 300 trials per session. For this experiment, the trials were interleaved and their order randomised.

### Spike sorting

We concatenated all AP band (bandwidth 300-10,000 Hz) binary files from a session for spike sorting, which we performed with Kilosort 2.5 (Pachitariu *et al*., 2024) and then manually curated with the Phy2 GUI. We classified clusters as putative single units if they exhibited (1) large spike amplitudes relative to baseline noise, (2) waveforms that were distinct from neighbouring clusters, and (3) a clear refractory period (>1 ms) evident in the autocorrelogram. We only included single units in subsequent analyses and excluded all action potentials from other clusters.

### Laminar and cell-type identification

We determined the laminar locations of individual S1 units and LFP channels based on the spatial distribution of waveforms along the probe, which we registered to physiological indicators of cortical layers, as described and validated previously (Emanuel *et al*., 2021). We defined the centre of layer IV as the depth corresponding to the earliest current sink in the local field potential current source density (CSD) at the onset of skin indentation. To generate CSD plots, we low-pass filtered voltage waveforms from the LFP band (0.5-500 Hz) at 250 Hz with a third-order Butterworth filter and then calculated the second derivative of this signal across laminar locations on the probe. We rigidly adjusted the depth of S1 units and LFP channels so that the centre of layer IV identified in the CSD plot was assigned to 476 μm below the cortical surface. S1 units and LFP channels were classified into cortical layers according to the following layer depths: layer II/III: 119–416.5 μm; layer IV: 416.5–535.5 μm; layer V: 535.5–952 μm; layer VI: 952-1200 μm. We classified S1 units as regular-spiking (RS) or fast-spiking (FS) based on the trough-to-peak times of the spike waveform on the channel with the largest amplitude (Bartho *et al*., 2004; Niell & Stryker, 2008). RS units were designated as those with a trough-to-peak time >0.55 ms and FS units were designated as those with a trough-to-peak time ≤0.55 ms.

### Analysis of optically evoked spiking activity

For optical stimulation sessions, we calculated z-scored firing rates in 20-ms bins using the mean and standard deviation in the baseline time window (0.8 s before the first laser pulse onset in each trial). We used a parameter-free ZETA test (Montijn *et al*., 2021) to identify whether each unit showed a time-locked modulation of spiking activity in response to optical stimulation with a p-value cutoff of 0.05. To identify sustained responses, we applied the ZETA test to the sustained time window (200-500 ms) for certain stimulus patterns. We compared the onset (0-60 ms) and sustained (200-500 ms) activity in synchronous and asynchronous conditions with the Wilcoxon signed-rank test. We estimated the optical receptive fields of S1 units by calculating the number of pedal pads that, when illuminated, produced S1 responses that exceeded a z-scored firing rate criterion, which we varied between 0.1 and 3.0, in any bin between 0.0 and 0.5 s after pulse-train onset.

### Analysis of local field potential response to optical stimulation

We aligned LFP waveforms to the onset of optical stimulation of each trial and computed the mean LFP waveforms for each pad for synchronous and asynchronous stimulation conditions. For each recording session, we averaged LFP waveforms from the same layer (layer II/III, layer IV, and layer V), normalised them by dividing the maximum response magnitude of the layer IV LFP, and then measured the magnitude of normalised waveforms during the stimulus onset window (0-60 ms) and sustained window (200-500 ms). We compared these between synchronous and asynchronous conditions using the Wilcoxon signed-rank test.

## Results

### Activation of the opsin CatCh generates high-frequency firing in Aβ LTMRs

For both mouse lines (*TrkC^CreER^;Avil^FlpO^;R26^LSL-FSF-Catch^* and *TrkB^CreER^;Avil^FlpO^;R26^LSL-FSF-Catch^*), only axons of the large-diameter sensory neurons of interest were labelled within forepaw glabrous skin (Fig. 1A). We first validated that the kinetics of the CatCh opsin were sufficiently fast to enable Aβ LTMRs to follow high-frequency trains of laser stimulation. To do so, we recorded from Aβ LTMRs in L4 DRGs with multiunit electrode arrays while optically stimulating the glabrous skin of the mouse’s ipsilateral hindpaw. We mapped receptive fields by directing pulse trains of 10 spikes in 0.25 s (rate of 40 Hz) to random spots on a grid centred on the suspected receptive field hot spot (example in Fig. 1B). When the stimulus was directed to the most distal part of the optical receptive field, which corresponded to the location of the mechanical receptive field, action potentials followed the laser patterns reliably. Less reliable responses were achieved when the laser was directed to more proximal sites (likely activating the axon within the nerve rather than at its terminal) in that an action potential only fired in response to the first laser pulse.

**Figure 1.**
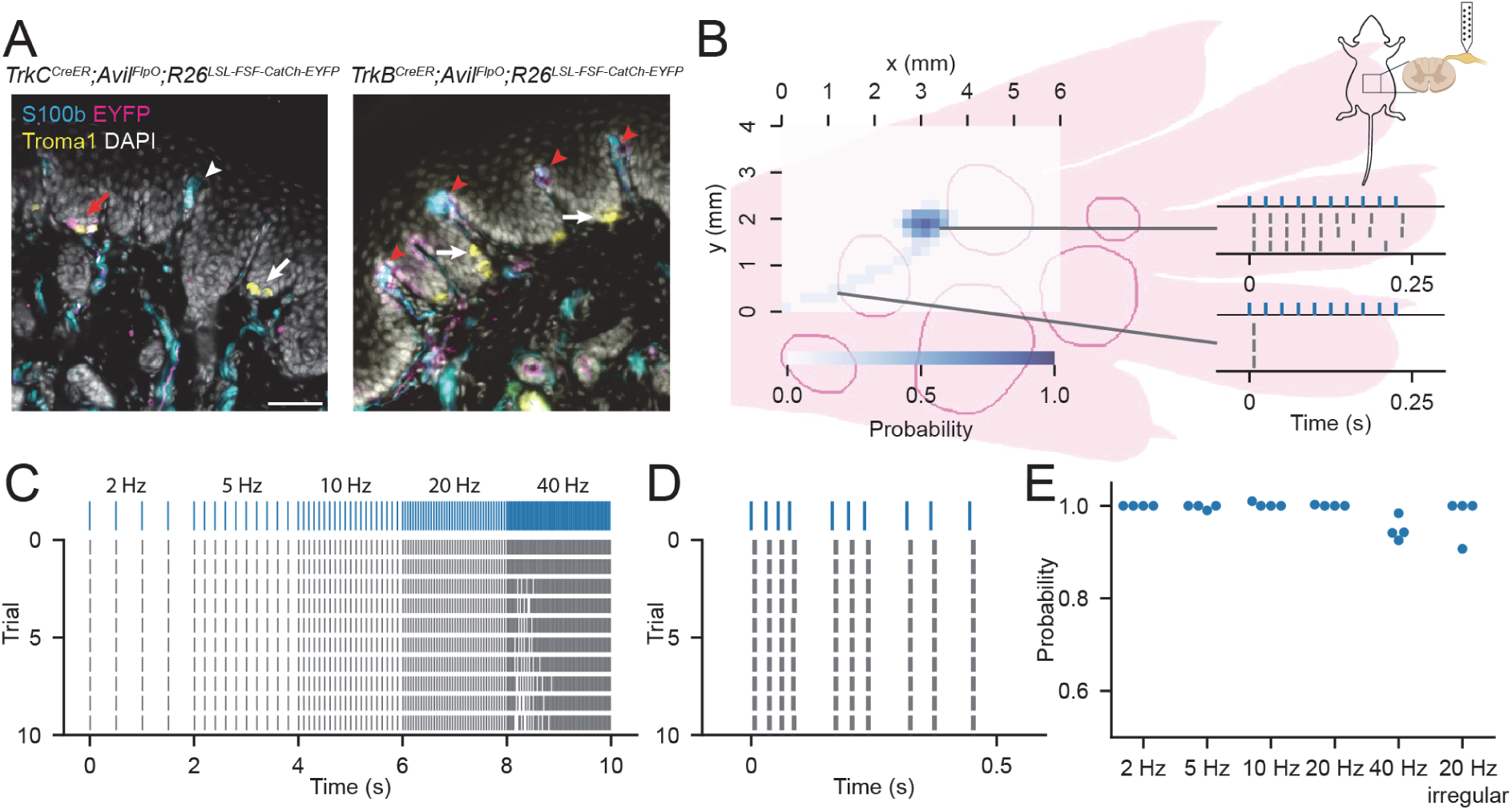
Optical Responses in Aβ LTMRs. (A) Aβ LTMR subtypes selectively labelled in immunostained forepaw sections of a *TrkC^CreER^;Avil^FlpO^;R26^LSL-FSF-Catch-EYFP^* mouse (left) and a *TrkB^CreER^;Avil^FlpO^;R26^LSL-FSF-Catch-EYFP^* mouse (right). Scale bar: 50 μm. Arrows indicate Troma1+ Merkel cells (head and shaft) and S100+ Meissner corpuscles (head only) that were associated with GFP+ (GFP antibody binds EYFP) fibres (red) or not associated with GFP+ fibres (white). (B) The optical receptive field of an Aβ RAI-LTMR measured by delivering 10-pulse trains (at 40 Hz) to multiple locations on the glabrous skin of a mouse’s hindpaw. The colour indicates the probability of laser-evoked action potentials. Raster plots show action potentials generated by the stimulus when directed at the axon terminal (top) or the nerve (bottom). (C) Raster plot showing action potentials generated by a series of laser pulses in ascending frequencies (2-40 Hz). (D) As in (C) for a train of 10 laser pulses with variable interpulse intervals (mean pulse rate of ∼20 Hz). (E) The probability of evoked action potentials for each stimulus in all four Aβ RAI-LTMRs.

Then, we stimulated the hot spot of the receptive field with trains of laser pulses. In the example neuron, the probability of action potentials after each laser pulse was 100% for trains of 2-20 Hz pulses and was 92.5% for trains of 40 Hz pulses (Fig. 1C). The probability of action potentials after each laser pulse was also 100% for a train of pulses at 20 Hz with irregular pulse intervals (Fig. 1D). Similar responses were evoked in all CatCh-expressing Aβ LTMRs tested. The latency of responses to laser pulses was between 4.75-6.80 ms, which is consistent with latencies measured after optical activation of ReaChR-expressing Aβ LTMRs (Emanuel *et al*., 2021). The probability of action potentials after each laser pulse was nearly 100% for 2-20 Hz pulse trains and greater than 90% for the 40 Hz and the irregular-pulse train (Fig. 1E). This demonstrates that CatCh expression enables LTMRs to follow pulse trains at physiological frequencies up to 40 Hz.

### Selective activation of Aβ SAI-LTMRs drives S1 neurons

We next examined how S1 neurons respond to optogenetically-evoked spike trains in Aβ SAI-LTMRs. In *TrkC^CreER^;Avil^FlpO^;R26^LSL-FSF-CatCh^* mice (n = 2) that expressed the opsin CatCh in Aβ SAI-LTMRs, optical stimulation directed to the centre of the forepaw generated robust S1 responses, consistent with the S1 responses generated with optogenetic activation of single action potentials in ReaChR-expressing Aβ SAI-LTMRs (Emanuel *et al*., 2021). Light-responsive units (n = 112 from 6 recording sessions) are shown in Fig. 2. At a low stimulation frequency (5 Hz), units generated a response to each laser pulse. At higher frequencies (20 Hz and 40 Hz), the spike patterns in the sustained phase of 57% of the units (64 of 112) were no longer distinguishable from baseline periods, indicating that they fully adapted. A similar pattern was observed in response to a pulse train built from a recorded spiking response of a TrkC^+^ Aβ SAI-LTMR during a 0.5-s, 20 mN skin indentation (data from Emanuel *et al*., 2021); the spiking pattern during the sustained phase differed from that at baseline in 31 of 112 units (27.7%).

**Figure 2.**
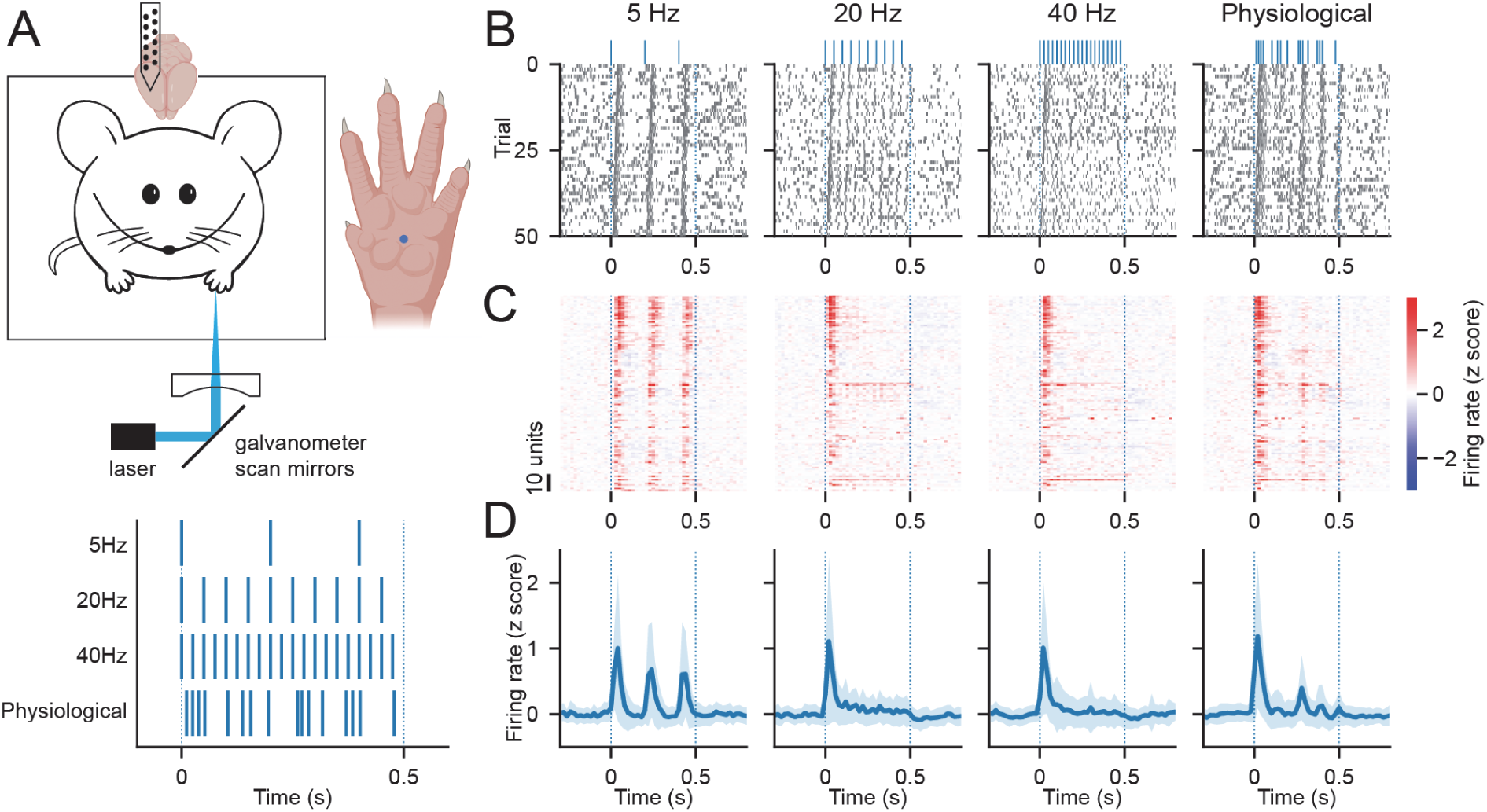
S1 responses to selective activation of Aβ SAI-LTMRs. (A) Top: An illustration of the optical stimulation and S1 recording setup. Bottom: Temporal patterns of optical stimulation, which was delivered to the centre of the left forepaw. (B) Raster plots of one example S1 unit in response to each stimulus pattern. (C) Heatmaps of z-scored firing rate responses of S1 units during optical stimulation at the paw centre with each pattern. (D) Z-scored firing rates (mean ± SEM) of S1 units (n = 112 units from 6 sessions).

These response patterns are dissimilar to those evoked by mechanical stimuli; activity during the sustained phase is extremely rarely observed in S1 in response to 20-mN indentations (Emanuel *et al*., 2021). One potential explanation for this difference is that the pulsed light activates multiple Aβ SAI-LTMRs synchronously whereas static indentation generates irregular spiking patterns in Aβ SAI-LTMRs (Iggo & Muir, 1969; Wellnitz *et al*., 2010). The irregularity of Aβ SAI-LTMR spiking during static indentation likely results in desynchronisation of their population-level firing pattern. Our artificially imposed synchrony may generate signals that propagate to S1, including during the sustained period. Sustained responses are exceedingly rarely generated by controlled-force indentations of intensities up to 20 mN (Emanuel *et al*., 2021).

### Synchronous activation of Aβ LTMRs generates sustained S1 activity

Therefore, we next directly tested whether the synchronicity of activation of Aβ SAI-LTMRs governs the propagation of their signals to S1, using another cohort of *TrkC^CreER^;Avil^FlpO^;R26^LSL-FSF-CatCh^* mice (n = 6). For comparison, we applied the same stimuli to *TrkB^CreER^;Avil^FlpO^;R26^LSL-FSF-CatCh^* mice (n = 7) to measure S1 unit responses to synchronous and asynchronous optical stimulation of Aβ RAI-LTMRs. We designed optical stimulation patterns to emulate synchronous or asynchronous recruitment of Aβ SAI-LTMRs with irregular pulses at mean rates of 20 Hz, which corresponds to the sustained firing rate of Aβ SA-LTMRs at moderate forces (∼20 mN; Emanuel *et al*., 2021). On each trial, we delivered laser pulses to five discrete locations within one of five pedal pads of the mouse’s forepaw either synchronously or asynchronously (Fig. 3A).

**Figure 3.**
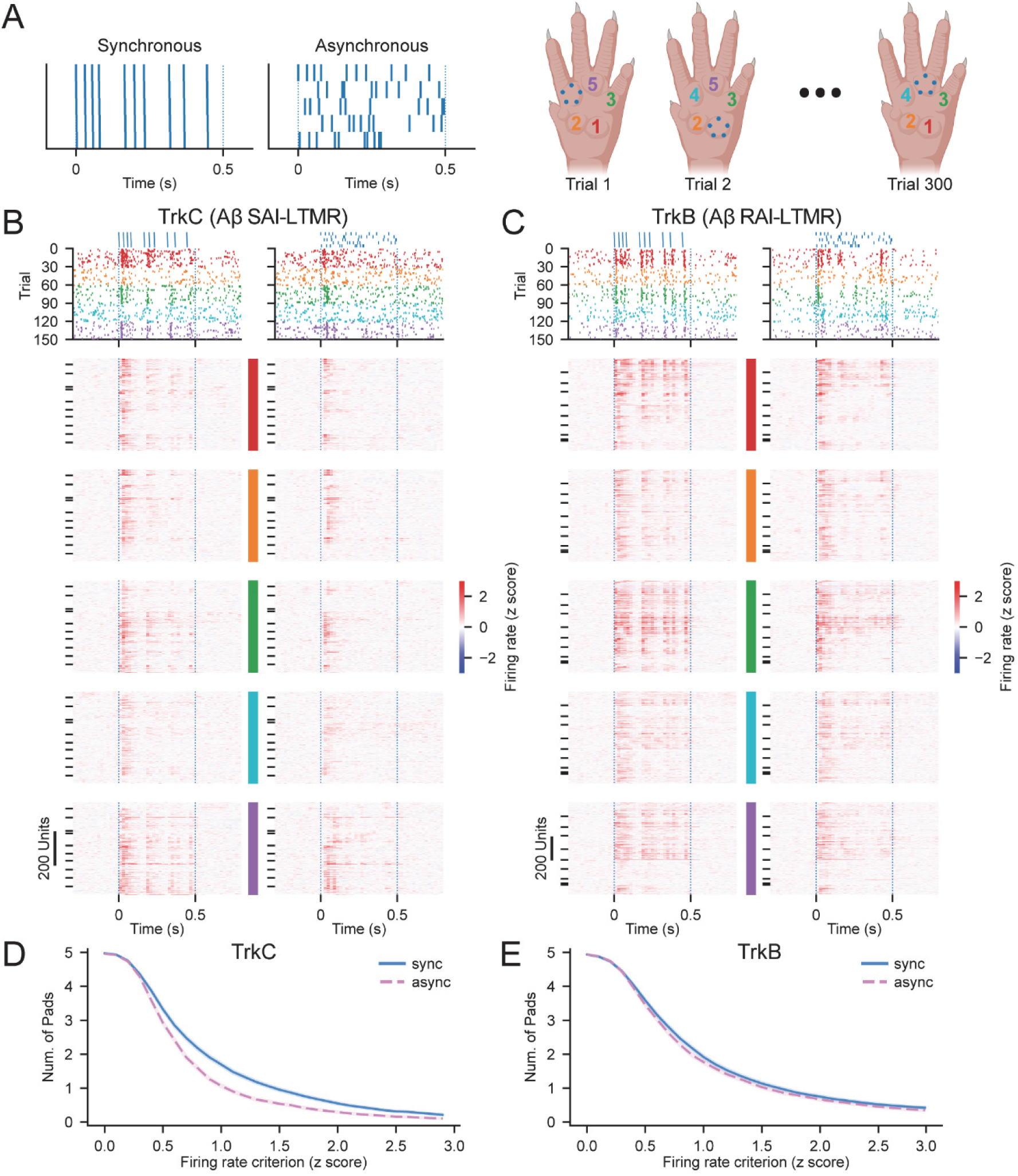
Activation of Aβ SAI-LTMRs and Aβ RAI-LTMRs at different forepaw locations with synchronous and asynchronous generates heterogeneous S1 responses. (A) On each trial, synchronous or asynchronous stimulation patterns were delivered to five different locations on one of five forepaw pedal pads. (B) Raster plot example (top) and heatmaps (bottom) of S1 unit responses to optical stimulation of Aβ SAI-LTMRs (TrkC) with synchronous and asynchronous patterns at each of the five pads on the mouse left forepaw. Units grouped by recording sessions. Within each recording session, units sorted by depth from the cortex surface. (C) As in (B) for responses to optical stimulation of Aβ RAI-LTMRs (TrkB). (D) Number of pads (mean ± SEM) that recruited a response in S1 units (n = 539) as a function of the maximum firing rate criterion (z scored) for synchronous and asynchronous stimulation of Aβ SAI-LTMRs. (E) As in (D), for stimulation of Aβ RAI-LTMRs (n = 773 S1 units).

Many S1 units (45.6%; 539 of 1183 units collected in 11 recording sessions) responded to optical activation of Aβ SAI-LTMRs with a change in firing pattern, typically an increase in firing rate (Fig. 3B). A larger fraction (60.3%; 773 of 1282 units collected in 10 recording sessions) responded to optical activation of Aβ RAI-LTMRs (Fig. 3C), consistent with observations made using ReaChR (Emanuel *et al*., 2021). For any individual unit, the response to each of the stimuli varied considerably. Notably, the number of pads that recruited a response after activation of Aβ SAI-LTMRs was greater with synchronous stimulation than with asynchronous stimulation (Fig. 3D; 1.70 ± 1.31 [mean ± SD] and 1.07 ± 1.06 pads for a criterion of max(z)>1, respectively; W = 5234, p < 0.001, n = 539 units, Wilcoxon signed-rank test), which suggests that the size of S1 unit optical receptive fields depend on how synchronously Aβ SAI-LTMRs are activated. Synchronous activation of Aβ RAI-LTMRs also recruited responses from more pads than asynchronous stimulation of the same neurons (1.92 ± 1.54 and 1.77 ± 1.47 pads, respectively; W= 32248.5, p < 0.001, n = 773 units), though the difference was much smaller in magnitude, even when comparing across firing rate criterion (Fig. 3E).

When considering responses to stimulation of the best pad, synchronous stimulation systematically produced larger magnitude onset (0 to 60 ms after first pulse) and sustained (200 to 500 ms after first pulse) responses than asynchronous stimulation of Aβ SAI-LTMRs (Fig. 4; ON: W = 26592.5, p < 0.001; sustained: W = 20100, p < 0.001, n = 539 units), but only of sustained responses with activation of Aβ RAI-LTMRs (onset: W = 139280.5, p = 0.289; sustained: W = 39836.5, p < 0.001, n = 773 units). This pattern was similar in both FS and RS units (Fig. 4C).

**Figure 4.**
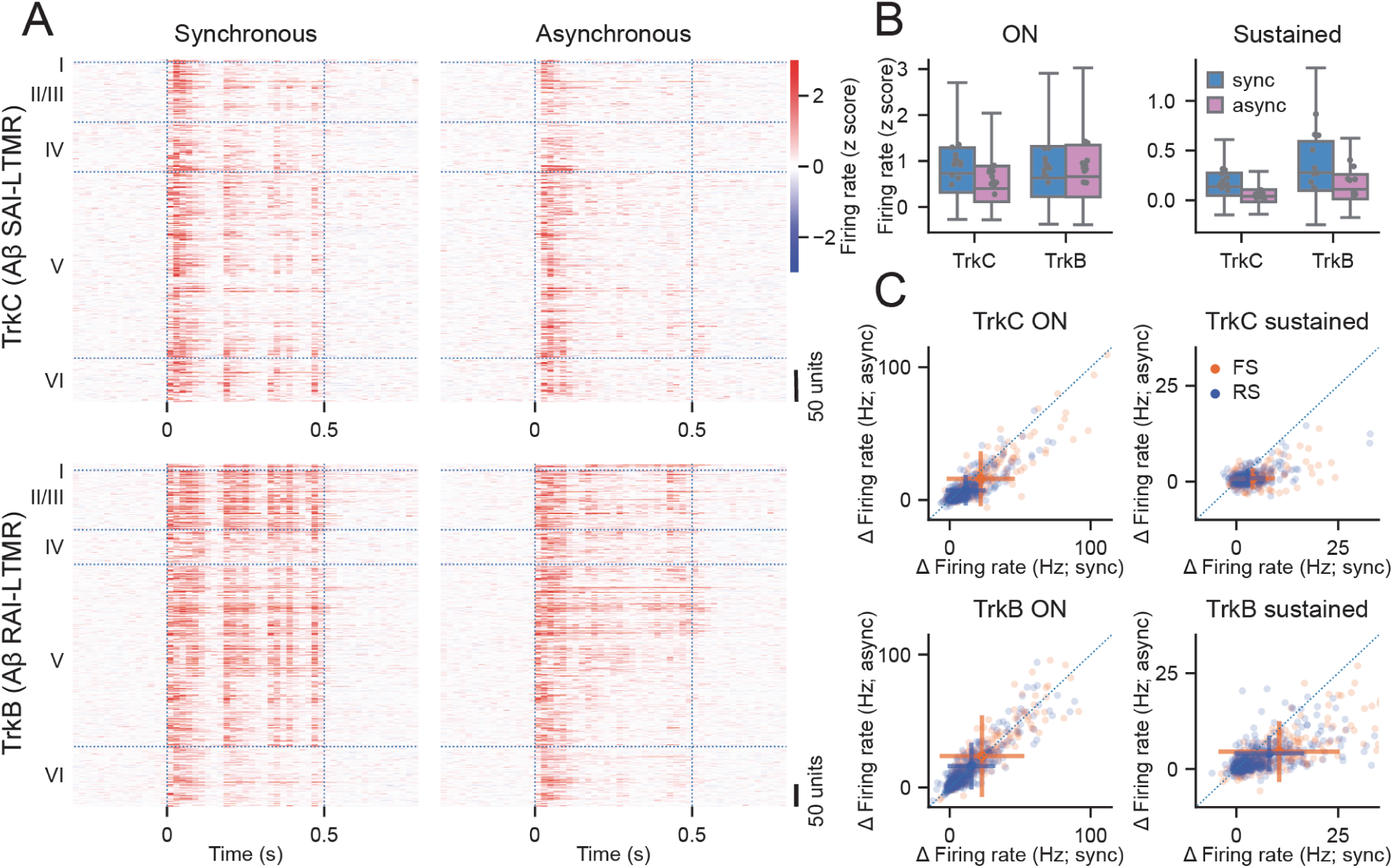
S1 responses to synchronous and asynchronous activation of Aβ SAI-LTMRs and Aβ RAI-LTMRs. (A) Heatmaps of S1 units in response to optical stimulation of Aβ SAI-LTMRs (TrkC) or Aβ RAI-LTMRs (TrkB) with synchronous and asynchronous patterns. Each row represents the z-score firing rate of one unit when synchronous and asynchronous stimuli were applied to the most responsive pad. Units sorted by their depth from the cortex surface. (B) Onset and sustained z-scored firing rate during synchronous (sync) and asynchronous (async) stimulation. The boxes show the quartiles of the z-scored firing rates while the whiskers extend to 1.5 inter-quartile range of the lower and upper quartile. Points represent mean response in individual recording sessions. (C) Scatterplots showing the baseline-subtracted firing rate onset (left) and sustained (right) responses of individual units for synchronous and asynchronous activation of Aβ SAI-LTMRs (top) or Aβ RAI-LTMRs (bottom). The large points and error bars indicate mean ± SD firing rates of all FS or RS units. Number of units: TrkC: n = 194 FS; 345 RS. TrkB: n = 257 FS; 516 RS.

### Synchronous activation of Aβ SAI-LTMRs generates stronger sustained local field potentials in S1

To evaluate the summed activity of S1 neuron populations, we extracted the local field potentials (LFPs) during synchronous and asynchronous activation of Aβ LTMRs. Trial-averaged LFPs revealed layer-specific patterned responses to optical activation of either cell type (Fig. 5A). To summarise across recordings, we averaged LFP responses from the pedal pad with the largest responses across channels from each cortical layer and normalised to the maximum response in layer IV (Fig. 5B). Synchronous activation of Aβ SAI-LTMRs generated stronger sustained activity than asynchronous activation for layer IV and layer II/III (layer IV: W = 8, p = 0.024; layer II/III: W = 9, p = 0.032, n = 11, Wilcoxon signed-rank tests) but not for layer V (W = 27, p = 0.638). Synchronous activation of Aβ RAI-LTMRs also generated sustained responses, but the response magnitudes did not significantly differ from asynchronous activation (layer II/III: W = 18, p = 0.375; layer IV: W = 18, p = 0.375; layer V: W = 18, p = 0.375, n = 10).

**Figure 5.**
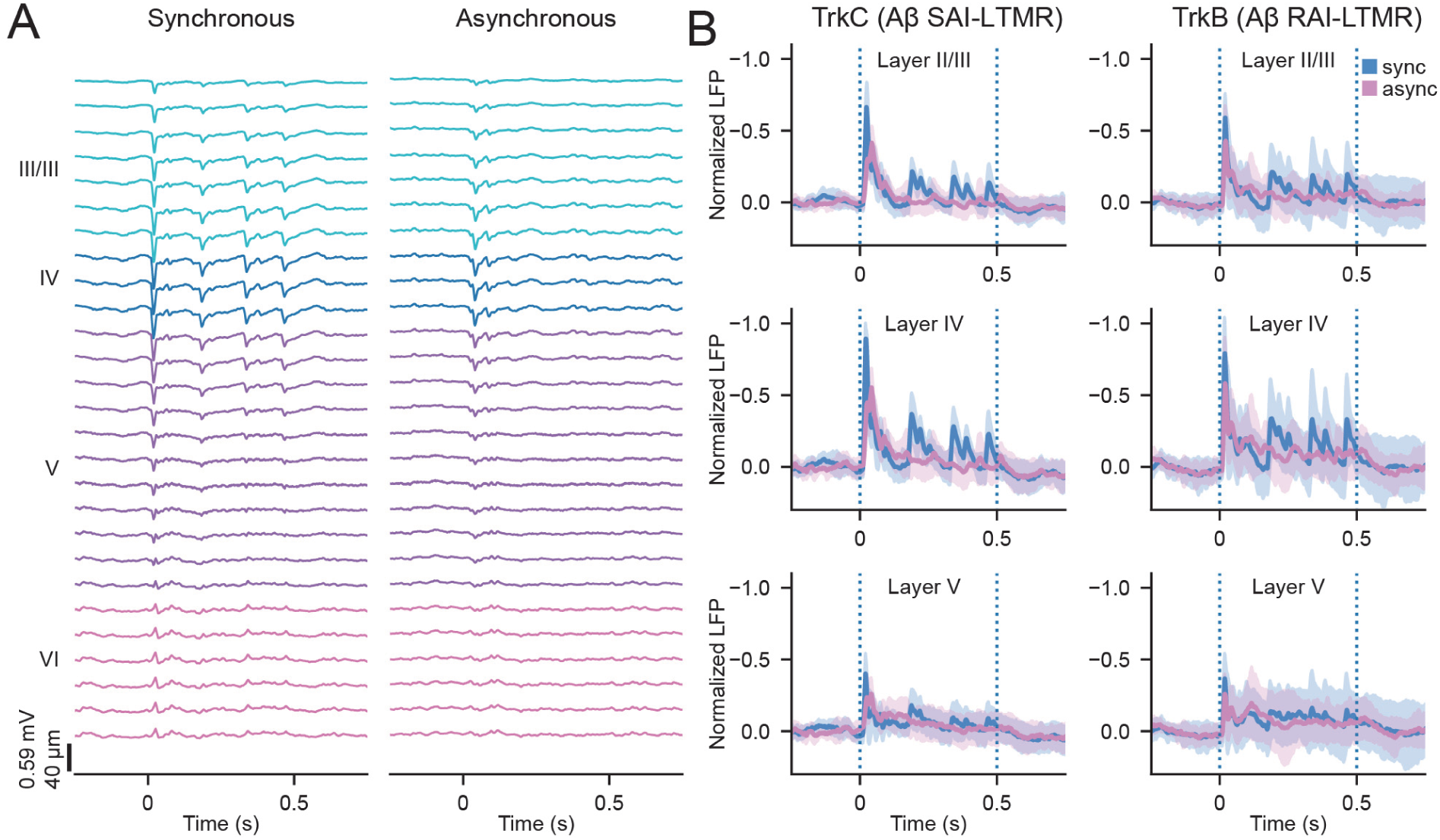
S1 local field potentials during synchronous and asynchronous activation of Aβ LTMR subtypes. (A) Trial-averaged LFPs during synchronous and asynchronous activation of Aβ SAI-LTMRs across S1 layers from one recording session. (B) The normalised LFP (mean ± SD) during synchronous and asynchronous activation of Aβ SAI-LTMRs (TrkC; left; n = 11 sessions) and Aβ RAI-LTMRs (TrkB; right; n = 10 sessions).

## Discussion

In this study, we used an opsin with fast kinetics (CatCh) to assess how selective activation of Aβ LTMR subtypes at physiological rates contributes to S1 responses. Using *in vivo* DRG recordings, we validated that pulsed optical stimulation of the axon terminals of CatCh-expressing Aβ LTMRs can reliably produce action potentials, even at rates as high as 40 Hz. Then, we investigated how stimulation of Aβ LTMR subtypes with distinct temporal patterns contributes to the S1 cortical response. We found that selectively activating either Aβ SAI-LTMRs or Aβ RAI-LTMRs generates responses in many S1 neurons, consistent with previous observations (Emanuel *et al*., 2021). Unexpectedly, activation of Aβ SAI-LTMRs with physiological temporal patterns generated prolonged increased firing in a subset of S1 neurons, perhaps because stimulation in the centre of the paw tended to recruit multiple Aβ SAI-LTMRs synchronously. Synchronous activation of Aβ SAI-LTMRs and Aβ RAI-LTMRs generated stronger responses in S1 than asynchronous activation. When the respective Aβ LTMR subtype was synchronously activated, S1 neurons showed more sustained activity compared to when those LTMRs were asynchronously activated.

While selective activation of Aβ SAI-LTMRs or of Aβ RAI-LTMRs generated responses in S1, their response properties were not identical. Selective activation of Aβ RAI-LTMRs produced stronger responses in S1. Furthermore, while activation of Aβ SAI-LTMRs or Aβ RAI-LTMRs both generate reliable onset responses in S1, activation of Aβ RAI-LTMRs generates stronger sustained activity that showed larger variability between recording sessions. Despite the subcortical convergence and integration of inputs from Aβ LTMR subtypes (Emanuel *et al*., 2021; Suresh *et al*., 2021; Chirila *et al*., 2022), Aβ RAI-LTMRs and Aβ SAI-LTMRs appear to differentially recruit subcortical circuits that regulate S1 response properties.

The characterisation of DRG responses revealed that functional CatCh was expressed along the axon fibres of sensory neurons. Thus, optical stimulation of the skin will activate the axon terminal as well as some fibres. However, only the axon terminal consistently followed high frequency laser pulse trains. Therefore, stimulating Aβ LTMRs with high frequency laser pulses principally recruits the axon terminals of the LTMRs. We also observed heterogeneous optical responses in S1. This response variability likely arose due to stochastic labelling of Aβ LTMRs as well as the variability in the precise location of the recording electrode relative to the S1 somatotopic map. Our asynchronous and synchronous stimulation patterns were targeted to five spots within each pedal pad, without knowledge of the actual location of CatCh-expressing axon terminals. This, along with the propensity of skin to scatter light, makes it difficult to estimate the number of neurons recruited with each laser pulse. Nevertheless, the differential effects of synchronous versus asynchronous activation across of both subtypes indicate that the spatial pattern of illumination recruited multiple distinct neurons during the sustained phase of stimulation.

Our results demonstrate that synchronous activation of Aβ SAI-LTMRs or Aβ RAI-LTMRs generates signals that propagate through the ascending pathways and drive cortex more effectively than asynchronous activation. This is consistent with predictions from neural network models, in which synchronous and asynchronous action potentials regulate the propagation of signals and multiplex rate coding and temporal coding in the central nervous system (Kumar *et al*., 2008; Ratte *et al*., 2013). Synchronous spiking facilitates stable signal propagation through neural networks (Kumar *et al*., 2008; Zhou *et al*., 2010) whereas asynchronous spiking and its associated variability can stabilise communication and increase information coding (Padmanabhan & Urban, 2010; Gast *et al*., 2024; Terada & Toyoizumi, 2024).

Distinct stimuli are likely to differentially drive synchronous and asynchronous firing in Aβ LTMRs. As noted above, irregular spiking patterns are signatures of Aβ SAI-LTMR static responses and are expected to produce asynchronous spiking patterns during sustained indentations. However, Aβ SAI-LTMRs produce larger responses with more regularity during dynamic stimuli (Iggo & Muir, 1969), which are likely to be synchronised across Aβ SAI-LTMRs with nearby receptive fields. The resultant enhanced propagation may allow for efficient cortical encoding of stimulus shapes and contours with high spatial contrast. A similar multiplexed representation has been observed in the olfactory bulb and S1 (Friedrich *et al*., 2004; Lankarany *et al*., 2019). With multiplexing of Aβ SAI-LTMRs signals, different components of the response could be used to influence separate downstream processes. Salient, high-contrast features from dynamic sampling may efficiently propagate through the system for perception through synchronous spiking. Maintained stimulus contact, however, that is encoded through asynchronous action potentials may be used in the spinal cord or the brainstem to modulate signals from other sensory neurons to, for example, regulate pain (Gautam *et al*., 2024).

Enhanced synchrony in the peripheral somatosensory system is associated with nerve injury and pain perception (Devor & Wall, 1990; Kim *et al*., 2016; Zheng *et al*., 2022; Chen *et al*., 2023). The transmission of asynchronous spikes in dorsal column axons to the somatosensory cortex is blocked by feedforward inhibition and fails to produce paraesthesia (Sagalajev *et al*., 2024). Our results support the idea that synchronous peripheral neuron activation and convergence of multiple sensory neurons onto downstream targets facilitate propagation through the ascending somatosensory system. Developing ways to control the extent to which skin or peripheral nerve stimulation synchronises or desynchronises activity across sensory-neuron populations could enhance the capacity to modulate brain activity for therapeutics and better provide haptic feedback for prostheses (Graczyk *et al*., 2016; Valle *et al*., 2018; Pfeifer *et al*., 2021; Valle *et al*., 2024).

## Data Availability

Data will be deposited into a public repository upon manuscript acceptance.

## Competing Interests

The authors have no competing interests.

## Author Contributions

WL, AEW, and AJE conceptualised the project, WL and AEW collected and analysed data with guidance from AJE. All authors wrote the initial manuscript draft, contributed to editing, and approved the final submission.

## Funding

This project was supported by the National Institutes of Health (R00NS119739 to AJE).

## Acknowledgments

We thank Katherine Greenhut and Yuna Lee for assistance with genotyping and mouse husbandry, Ronghao Zhang for code for extracting local field potentials, and Christopher Rodgers and members of the Emanuel lab for discussions.

## Notes

### Competing Interest Statement

The authors have declared no competing interest.

